# Neural correlates of cognitive motor signals in primary somatosensory cortex

**DOI:** 10.1101/2020.04.20.041269

**Authors:** Matiar Jafari, Tyson NS Aflalo, Srinivas Chivukula, Spencer S Kellis, Michelle Armenta Salas, Sumner L Norman, Kelsie Pejsa, Charles Y. Liu, Richard A Andersen

**Author notes:** Contributed Equally (Matiar Jafari, Tyson NS Aflalo). **Contributions:** M.J., T.A, and R.A.A designed the study, M.J. and T.A developed experimental tasks, M.J. and T.A. designed analysis and analyzed the data, M.J., S.C., S.S.K, and M.A.S. collected the data, S.L.N contributed code, M.J. and T.A. interpreted results and wrote the original draft, M.J., T.A., S.C., and R.A.A. reviewed and edited the paper, T.A. and R.A.A. provided mentorship, T.A., S.K., and R.A.A. acquired funding, K.P provided administrative and regulatory assistance, C.Y.L. performed implantation surgery.

## Abstract

Classical systems neuroscience positions primary sensory areas as early feed-forward processing stations for refining incoming sensory information. This view may oversimplify their role given extensive bi-directional connectivity with multimodal cortical and subcortical regions. Here we show that single units in human primary somatosensory cortex encode imagined reaches centered on imagined limb positions in a cognitive motor task. This result suggests a broader role of primary somatosensory cortex in cortical function than previously demonstrated.

## Main

Somatosensory cortex (S1) is largely studied and understood in its role as the primary sensory region for processing somatic sensory signals from the body. However, recent work highlights a more direct role in motor production: S1 neurons can respond to passive movements alone, active movements alone, or both^1,2^ and neurons become activated prior to movement initiation^1,3^. S1 neurons project to the spinal cord^4,5^, and electrical or optical stimulation of S1 elicits motor movements^6–8^. These results indicate a direct role of S1 in the production of motor behavior. However, in many of these studies, it is hard to dissociate whether neural signals reflect motor variables or aspects of sensory processing.

To understand if S1 processes reach intentions in the complete absence of sensation or expected sensations, we recorded single unit activity from multi-channel arrays implanted in the sensory cortex (Fig 1B) of a 34-year-old tetraplegic male (FG) during a delayed imagined reaching task. The arrays were implanted in S1 as part of an ongoing clinical trial in which we showed that microstimulation delivered through these same multi-channel arrays evokes localized and naturalistic cutaneous and proprioceptive sensations^9^. Our paradigm (Fig 1A), adapted from previous non-human primate studies^10,11^, systematically manipulated fixation, imagined initial hand, and reach target locations at distinct points in the trial. Importantly, the subject is capable of moving his eyes and thus can direct his gaze to the fixation targets. However, the paralyzed subject did not move his arm, but instead used motor imagery to imagine moving his hand to the initial hand cue location and subsequently imagine moving it to the final target location. This design allowed us to: one, understand how activity in S1 relates to storing information about arm location, movement plans, and movement execution, and two, characterize the reference frame of these signals, e.g. whether movement variables are coded relative to the initial imagined position of the hand, relative to the eyes, or relative to the body or world.

**Figure 1.**
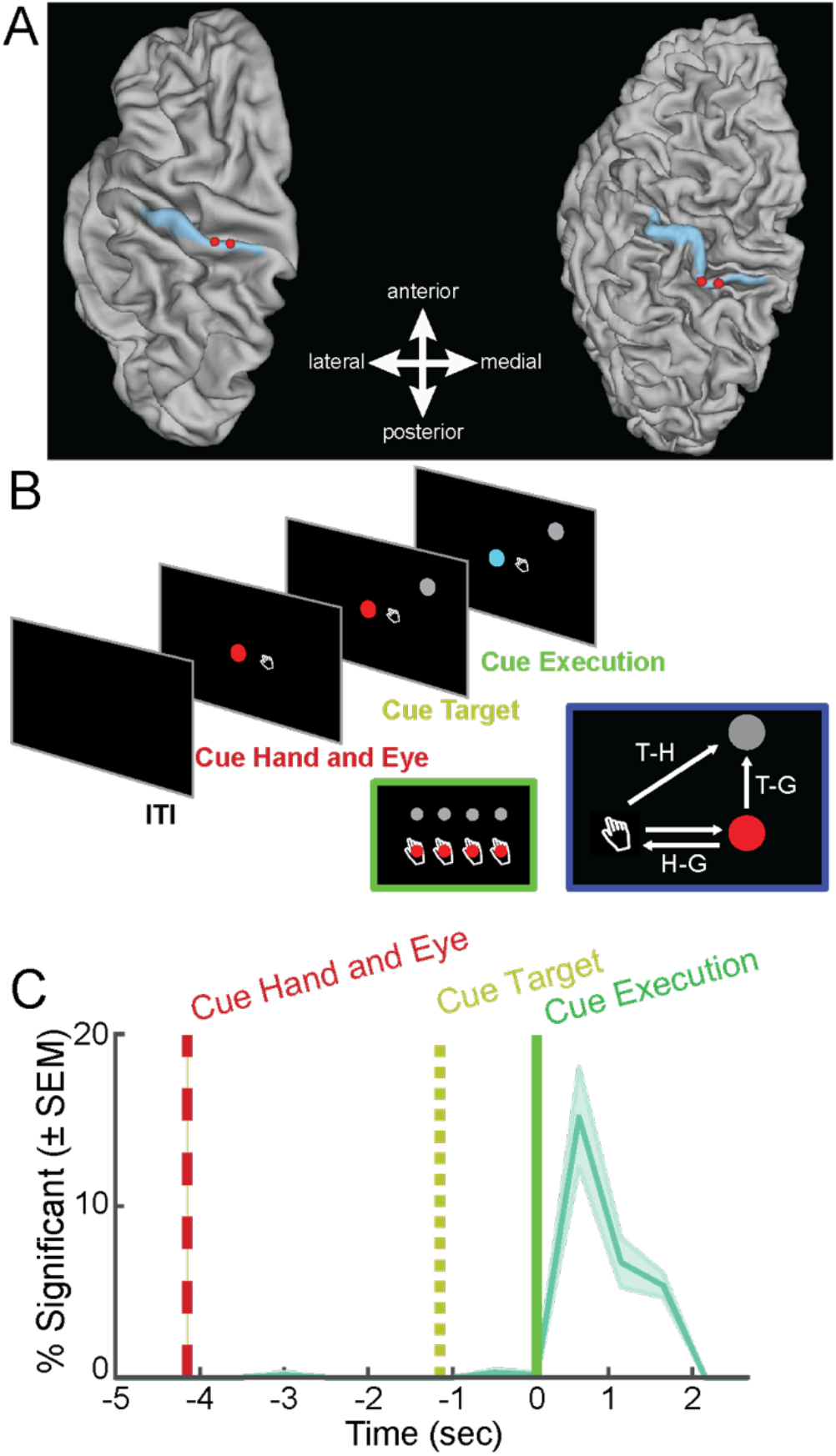
Behavioral Task, Electrode Array Location, and Tuning Throughout Task. **A)** Group-average brain map (left) and brain of subject FG (right) showing location of implanted microelectrode array (red) and Brodmann Area 1 (blue). **B)** Time course of delayed reference frame reaching task testing all unique combinations of four gaze, hand, and target positions (green inset). Geometry of the reference frame task (blue inset). **C)** Percent of units in population tuned (mean ± SEM, p < 0.05, FDR corrected). Units were considered tuned from the beta value of the linear fit, and subsequently corrected for multiple comparisons.

We performed a sliding window analysis to understand whether and when neurons in S1 become active for our cognitive motor task. For each unit, we used a linear model with interactions to explain firing rate as a function of fixation, initial imagined hand, and target locations (Fig 1C, p < 0.05 FDR corrected for number of units per time slice, window size: 750 ms, step size: 500 ms). We find negligible selectivity following cueing of the hand and eye positions indicating no neural coding for true eye position or the imagined position of the arm. We also found negligible selectivity following target presentation, indicating no encoding of the spatial location of the target or planning activity related to the upcoming imagined motor action. Finally, we found that a significant proportion of the population was selective following the instruction to initiate the imagined reach. Thus, sensory cortex is engaged during a cognitive motor task despite the absence of overt movement and sensory feedback, but only during imagined movement of the limb.

We found that nearly all the neurons selective during the movement execution phase coded movement as the reach vector: the direction of imagined movement of the hand. In other words, selective units coded the location of the target relative to the initial imagined hand position (or, by symmetry, hand position relative to the target). This result was found using a gradient analysis pioneered in NHPs^12^; neural responses for each unit were organized into response matrices where the firing rate is coded for each hand, eye, and target position. A gradient field is then computed which describes how the firing rate is sensitive to changes in the behavioral variables. Finally, the resultant, or vector sum, of the gradient field summarizes the net effect of behavioral manipulations which can be used to determine whether neural activity encodes target position relative to gaze position (T-G), the target position relative to the hand (T-H), the hand position relative to gaze direction (H-G), or some combination of these vectors (Fig S1). A representative response matrix for a neuron coding the position of the target relative to the hand and the population distribution of response gradients is shown in figure 2B and C. Interestingly, despite no neural coding for imagined hand position prior to movement execution, our result indicates that the neural coding during the imagined reach incorporates the internal estimate of hand position when computing the reach response.

**Figure 2.**
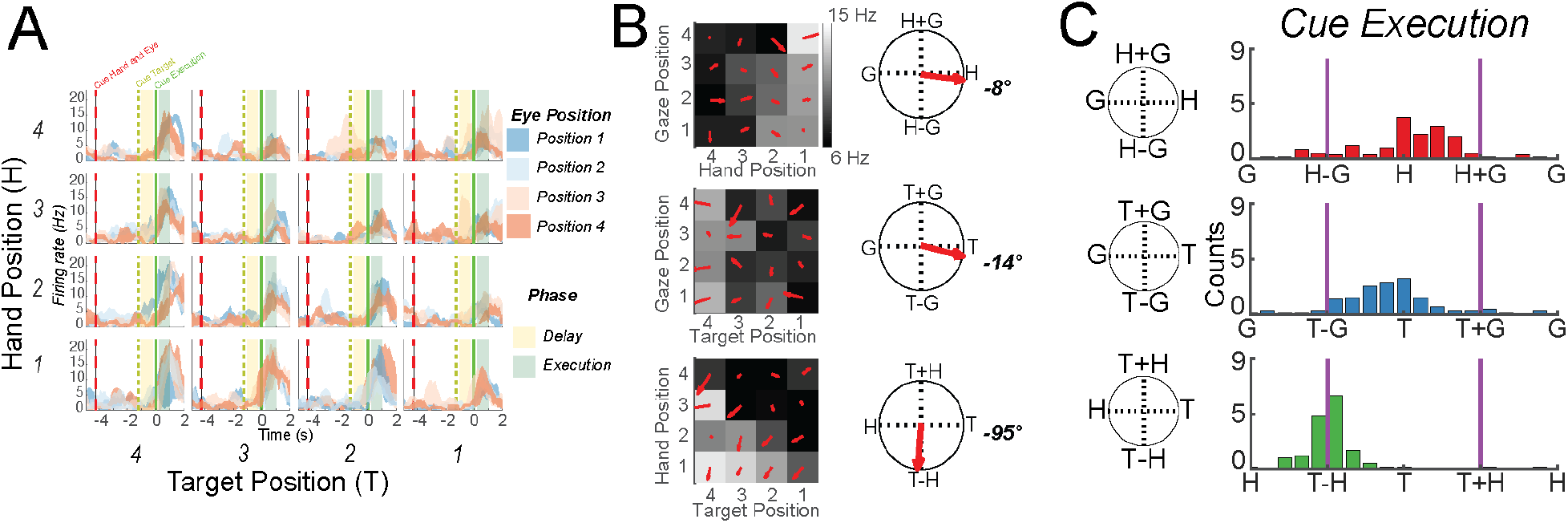
Example S1 Unit, Gradient Analysis, and Distribution of Gradients of Tuned Units. **A)** Peristimulus time histograms for all 64 conditions (3 trials; mean ± SEM). Each of the 16 subplots shows the response of the unit to a particular combination of eye, hand, and gaze position. **B)** matrices and gradient resultant orientations for the cell shown in 2A during the execution epoch (Fig 2A, green). **C)** Histograms show gradient resultant orientations for the population of tuned units.

Single unit analysis shows that the population is dominated by units coding the reach vector. To verify that this interpretation is an adequate summary of S1 encoding, we used complex principal component analysis (cPCA) to characterize the full temporal dynamics of the reference frame of the population as a whole (Fig S2). The gradient analysis described above summarizes the sensitivity of a neuron to behavioral variables using the resultant of the gradient, a 2D vector that can be described by a length and angle. We used cPCA for its capability to handle vector data samples, i.e. described by both a length and angle for each observation^13^. We find that coding of the reach vector strengthens and peaks around 750 ms after the cue to execute the imagined reach (Fig 3A). Further, only the first cPCA component was significant (parallel analysis^14^, α < 0.05). This suggests that reach coding in S1 is dominated by a single homogeneous representation of the reach vector that can be used to decode the subject’s motor intent (Fig 3B).

**Figure 3.**
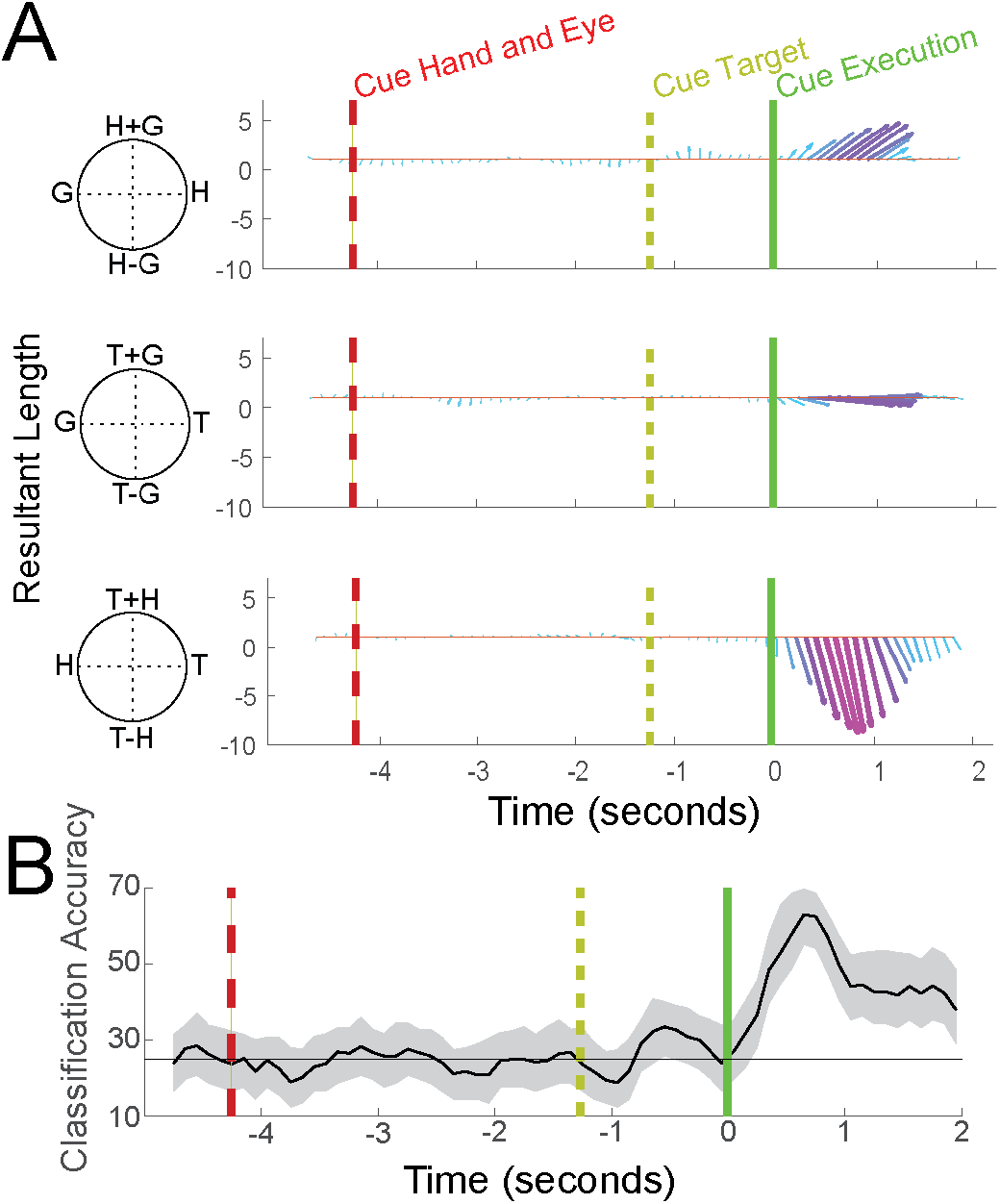
Population level reference frame coding and decode accuracy during a cognitive motor task in S1. **A)** Temporal evolution of reference frame encoding across the population of S1 units. Only the first component (shown) was found to be significant (p < 0.05; parallel analysis). Arrow length, width, and color shows tuning strength. **B)** Temporal dynamics of imagined reach decoding. Offline analysis depicting accuracy of target classification from the middle two hand positions [750-ms sliding window, mean with 95% confidence interval (CI)]. Horizontal black line is chance, 25%).

We have shown the first single unit evidence that cognitive imagery of movements is represented in the primary somatosensory area of human cortex. S1 neurons tracked motor execution intentions in the complete absence of sensation exclusively during imagined execution. There was negligible activity while the subject maintained the position of the limb in memory, fixated distinct targets, or planned movements. These results suggest that the role of S1 in motor production is restricted to the time of movement execution. Given the timing of S1 activity and reciprocal connections with motor cortex, S1 activity may reflect an efference copy of execution signals originating from motor cortex. Lastly, we show S1 activity codes motor intention relative to the imagined position of the hand. A possible concern is that these results are unique to individuals who have lost their main peripheral input. However, recent findings from our lab and others have shown these representations are largely stable and reorganization does not result in the production of novel functional sensory representations^9,15–17^ These results implicate a role of primary somatosensory cortex in cognitive imagery, a role of S1 in motor production in the absence of sensation, and suggest that S1 can provide control signals for future neural prosthetic systems.

## Acknowledgments

This work was supported by the National Institute of Health (R01EY015545, 5U01NS098975-02), the Tianqiao and Chrissy Chen Brain-machine Interface Center at Caltech, the Conte Center for Social Decision Making at Caltech (P50MH094258), the David Geffen Medical Scholarship, and the Boswell Foundation. The authors would also like to thank subject FG for participating in the studies, and Viktor Scherbatyuk for technical assistance.

## Data availability

All primary behavioral and neurophysiological data are archived in the Division of Biology and Biological Engineering at the California Institute of Technology and are available from the corresponding author on reasonable request.

## Code availability

All custom-written analysis code is available from the corresponding author on reasonable request.

## Materials and Methods

### Subject Information

Neural recordings were made from participant FG, a tetraplegic 32-year-old male with a complete C5/C6 spinal cord injury. FG was implanted 1.5 years post-injury for a clinical trial of a BMI system consisting of intracortical stimulation and recording. Neural recordings for the current study were acquired 1-year post implantation. All of subject FG’s sensations and motor ability are consistent with the level of the sustained injury. The subject remains intact for all other motor control and sensations above the level of injury. Surgical implantation took place at Keck Hospital of USC.

Experiments were conducted in the framework of an ongoing neural prosthetics clinical study (ClinicalTrials.gov identifier: NCT01964261) and were in compliance with all relevant clinical regulations. We obtained informed consent after explaining the objectives of the study and the possible risks involved. The study and all procedures were approved by the Institutional Review Boards (IRB) of the California Institute of Technology (Caltech), the University of Southern California (USC), and Rancho Los Amigos National Rehabilitation Hospital (RLA).

### Surgical Planning and Implantation

Surgical planning for subject FG followed the protocols described in^1,2^. In brief, functional magnetic resonance imaging (fMRI) was used to measure the BOLD response while FG performed imagined reaching and grasping movements in response to visual cues. The statistical parametric analysis guided selection of implant locations in the left hemisphere of the ventral portion of the premotor cortex (PMv), the supramarginal gyrus (SMG), and somatosensory cortex (S1). PMv and SMG were implanted with 96-channel Neuroport microelectrode arrays (Blackrock Microsystems, Salt Lake City, UT). S1 was implanted with two 7×7 microelectrode arrays (48 channels per array, Blackrock Microsystems, Salt Lake City, UT) on the post-central gyrus. Figure 1B shows the implantation locations for the two arrays. In addition, we estimated the anatomical location of the implantation of S1 in terms of Brodmann’s Area. To this end, we used Freesurfer^3^ to perform a surface reconstruction of the individual subject’s anatomy. The subject’s anatomy was then registered to the 164K fs-lr group-average template using Connectome Workbench^4^. The subject’s implants were determined to be localized to Brodmann’s Area 1 (BA1) according to the composite template of Van Essen et al 2012^5^ as visualized within Connectome Workbench. Localizing the areal boundaries of BA1 within the individual subject requires the registration of the individual subject’s surface anatomy to a group-average atlas. We therefore show implant locations both as they appear on the individual subject’s brain surface as well as where the arrays are estimated to be located on the fs-lr group-average template brain (Fig 1A).

### Reference Frame Task

Experimental sessions with subject FG were performed at Rancho Los Amigos National Rehabilitation Center (RLA). FG performed the task in a dimly lit room seated in his motorized wheel chair. Task stimuli were viewed on a 47-inch LCD monitor with the screen occupying approximately 45 degrees of visual angle. The subject was asked to minimize head movements throughout the task. At the beginning of each trial, FG was presented with a fixation cue and a hand position cue. Each cue could be positioned at one of four locations resulting in 16 possible hand and eye configurations. FG was able to move his eyes and thus fixate the fixation cue as verified using eye tracking. In contrast, FG did not position his actual hand at the location of the hand cue, but instead FG imagined moving his hand to the cue location and maintained imagery of his hand until the go cue. After 3 seconds a reach target cue was shown at one of four spatial locations arranged parallel to and above the cued eye and hand positions. The target cue was shown for 1.25 seconds during which the subject continued to hold their gaze and imagined hand positions. A change in the color of the fixation marker instructed the subject to begin an imagined reach to the cued target location. The subject was asked to make an imagined reach and maintain the imagined ending position (target location) until the execution epoch was over (2 seconds). The execution epoch was then followed by an inter-trial interval (ITI) of 2 seconds. A schematic representation of the task is shown in figure 1B.

Experimental data was collected in three experimental runs. Each run consisted of a total of 64 trials, one trial for each unique combination of the four eye, hand, and target positions. This resulted in 192 total trials, 3 repetitions for each unique trial type. Each experimental session was separated out by at least a week.

### Neural Recordings

Neural activity from each array was amplified, digitized, and recorded at 30 kHz using the Neuroport neural signal processor (NSP). The Neuroport system, comprised of the arrays and NSP, has received FDA clearance for less than 30 days of acute recordings. However, for purposes of this study we received FDA IDE clearance for extending the duration of the implant (IDE number: G130100).

Putative waveforms were detected at thresholds of −3.5 times the root-mean-square after high pass filtering the full bandwidth signal (sampled at 30Khz), using the Blackrock Central software suite (Blackrock Microsystems). Waveforms consisted of 48 samples, 10 prior to threshold crossing and 38 samples after. These recordings were then sorted (both single and multi-unit) using k-mediods clustering using the gap criteria to estimate the total number of clusters^6,7^. Offline sorting was then reviewed and adjusted as needed following standard practice^8^. On average across 4 days of recordings in S1 we analyzed 163 sorted units per session. All sorting was done prior to analysis and blind to channel or unit responses found during the task. Further spike sorting methods can be found in Zhang et al., 2017.

### Eye Tracking

Subject FG’s eye position was monitored using a 120 Hz binocular eye tracking system (Pupil Labs, Berlin, Germany). If the subject’s gaze shifted off the cued eye position the task was terminated and restarted to ensure that gaze position was correct and remained fixed to the cued eye position throughout each appropriate epoch for a run (64 consecutive trials). Eye positions were synced to the task and allowed online determination of eye position. We instructed the subject to maintain a constant head position and to only move his eyes to fixate the target. However, head position was not monitored and more conservatively we can say that we manipulated gaze as opposed to eye position proper. In either case, our results showed no dependences on eye/gaze and thus the distinction is not especially important given the pattern of results.

### Linear Analysis for Tuning (Fig 1C)

We defined a unit as selective if the unit displayed significant differential modulation for our task variables as determined by a linear regression analysis: We created a matrix that consisted of four indicator variables for each unique behavioral variable (e.g. one indicator variable for each of the four initial hand positions) resulting in 12 indicator variables. Firing rate was estimated as a linear combination of these indicator variables and their interactions: FR is firing rate, Xc is the vector indicator variable for condition c, βc is the estimated scalar weighting coefficient for condition c, and β0 is the intercept.

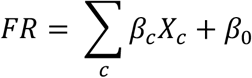

Linear analysis was performed over a sliding window throughout the interval of the task. Windows were 750 ms in duration and window start times were stepped every 500 ms. Significance of each fit was determined using the p-value of the F-test of overall significance for the linear analysis (p < 0.05, FDR-corrected for number of units). Units that were found to be significant in this analysis were then determined to be selective and further analyzed in the reference frame analysis.

### Reference Frame Analysis: Gradient Analysis (Fig 2)

Gradient analysis was used to quantify how changes in the behavioral variables changed the firing rate of each unit when comparing across each unique combination of variable pairs (Hand-Gaze (HG), Target-Gaze (TG), and Target-Hand (TH))^9–12^: For each tuned unit (based on the p-value of the linear regression model described above) we created a four by four matrix (response matrix) representing neural activity for each unique combination of two behavioral variables; thus, for example, the value at the HG response matrix location [x,y] would be the neural activity recorded for hand position x and gaze position y averaged across trial repetitions and repetitions acquired for the different target positions. Gradients were determined using the gradient function in Matlab 2019a (Mathworks Inc, Natick, MA). For each gradient, a resultant angle and length was computed to summarize the net direction and magnitude of change across the entire response matrix (Fig. S1). However, often times the gradients often show a symmetrical pattern that would result in cancellation of symmetrical angles (Fig. S1A). To avoid this, we double each angle in the matrix and represent each angle from 0° to ±180°. Therefore, the summed resultant angle is represented by 0° for gradients oriented left and right, ±180° for gradients oriented up and down, and −90° for gradients oriented along the diagonal (Fig S1A). The summed resultant angle and length however cannot be mapped directly onto the response matrix; thus, we have notated the appropriate variable and combinations of variables to help with interpretation. For example, in figure S1A hand only (H) modulation would be found at ±180°, gaze only (G) modulation is seen at 0°, H+G at 90°, and H-G at −90°. Therefore, we can use the angle of the resultant angle as a proxy for overall orientation bias for a variable or variable pair.

### Dimensionality Reduction (Fig 3)

We used population level dimensionality reduction analyses to determine the most common modes of reference frame encoding over time. This was done in a three stage process (see Fig S2): (1) Initial principal component analysis (PCA) on the time varying activity of the neural population, (2) reference frame analysis on each time point of the resulting principal components, (3) complex principal component analysis (cPCA) on the resultant angles and magnitudes. The initial principal component analysis was used to denoise and improve the calculation of reference frames at the level of the population. In order to perform PCA analysis we constructed a matrix of neural data D that was (n) by (t * c) in size, with n being the number of neurons, t being the number of time points, and c being the number of conditions. For each neuron, activity was averaged across repetitions of the same condition within a 100ms window. The reference frame analysis was then applied to each temporal window for the first 20 principal components. Note that following the initial principal component analysis, the population activity still carries detailed information about neural selectivity properties unrelated to the reference frame proper (e.g. preferred directions of movement or preferred hand locations.) Thus multiple principal components may have the same reference frame, but simply prefer a different direction movement direction. Computing the reference frame at this stage extracts the population level reference frame, abstracting away tuning preference differences. The final cPCA was then used to capture the main reference frame modes once the detailed aspects of tuning (e.g. such as preferred direction of response) were abstracted away by the reference frame analysis. We used cPCA given the fact that dimensionality reduction was performed on the resultant vectors, values with both a magnitude and angle. We converted all resultant angles and lengths into complex numbers to apply cPCA^13^. We used parallel analysis to determine which components from this dimensionality reduction were significant^14^.

### Discrete Classification (Fig 3B)

Offline classification was performed using linear discriminate analysis. The classifier took as input a vector comprised of the number of spikes occurring within a specified time epoch for each sorted unit. The following assumptions were made for the classification model: 1) the prior probability across the classes was uniform, 2) the conditional probability distribution of each feature on any given class was normal, 3) only the mean firing rates differ for each class (the covariance of the normal distributions were the same for each class), and, 4) the firing rates of each input are independent (covariance of the normal distribution was diagonal). Reported performance accuracy was based on leave-one out cross-validation. To compute the temporal dynamics of classification accuracy, the neural data was first aligned to a behavioral epoch (e.g. cue execution onset). Spike counts were then computed in 750 ms windows spaced at 100 ms intervals. Classification accuracy was computed independently for each time bin and bootstrapped resampling was used to compute 95% confidence bounds.

