## Supplemental figures for "Neural correlates of cognitive motor signals in primary somatosensory cortex"

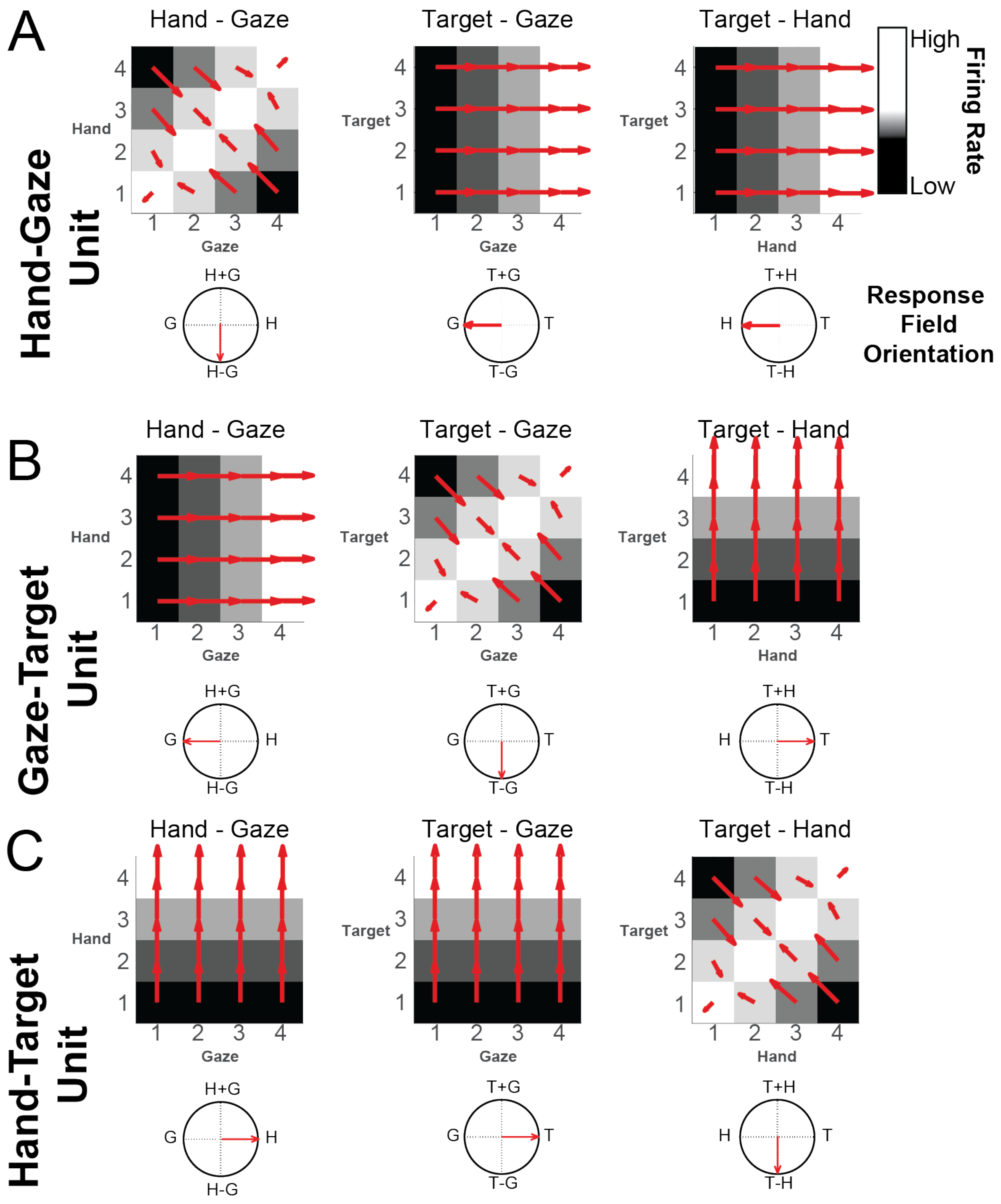


Figure S1. Idealized Unit Responses Displaying Eye-Hand-Target Tuning for Three Different Reference Frames. **A)** Hand-Gaze unit encoding the relative location of the hand and point of gaze. **B)** Gaze-Target unit encoding the relative location of the target and point of gaze. **C)** Hand-Target unit encoding the relative location of the hand and target. The response field orientations for all units are shown for each of the three variable pairs along the horizontal beneath each response matrix. Gradient analysis, and the subsequently derived resultant angle (±180º and 0º point left and right, respectively) and length, quantifies how firing rate responses can be attributed to each behavioral variable (HG, TG, and TH).

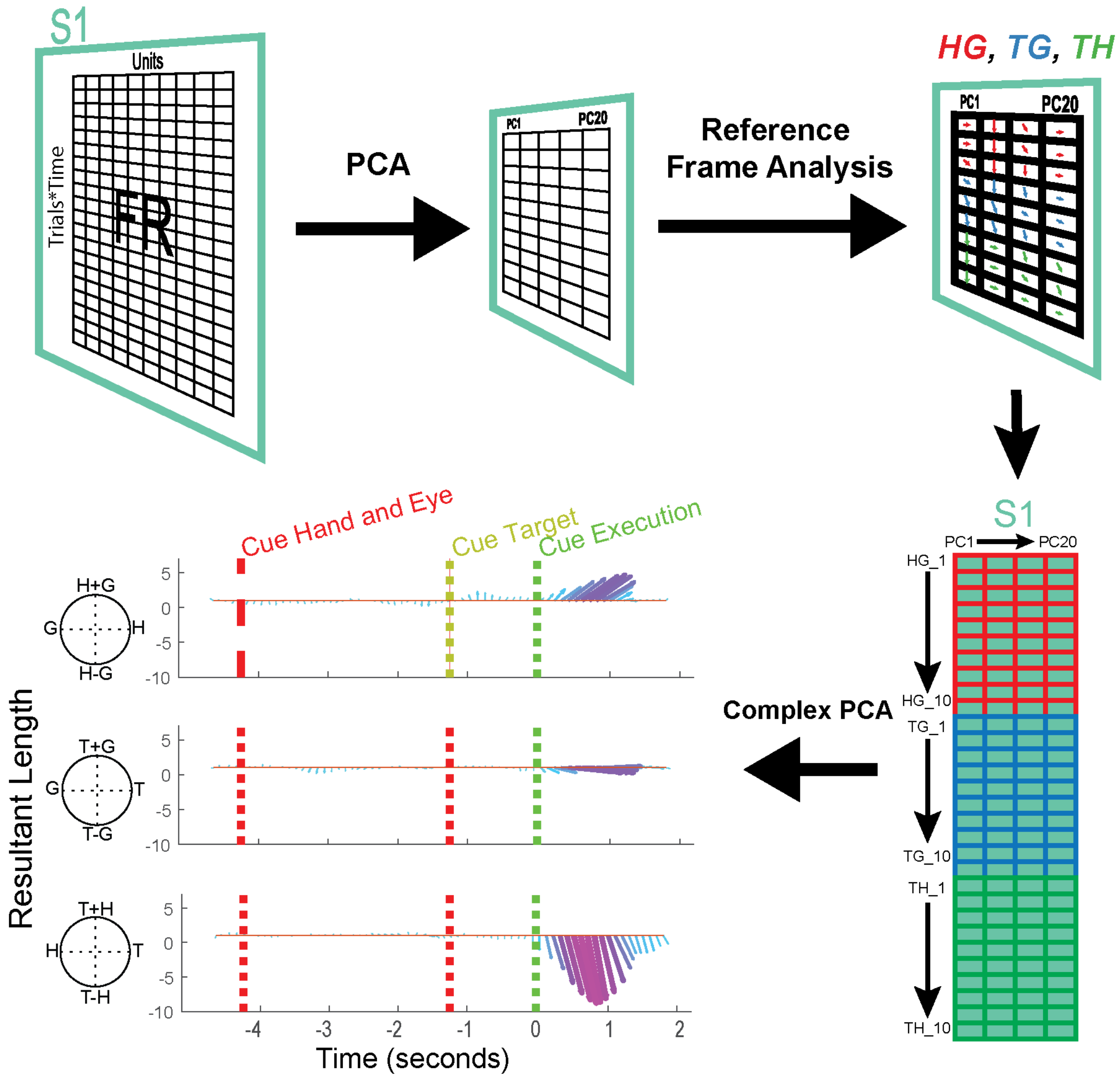


Figure S2. Graphical Illustration of Complex PCA Processing.
